# The Pik NLR pair accumulates at the plasma membrane as a hetero-oligomeric sensor-helper immune protein complex prior to activation

**DOI:** 10.64898/2025.11.30.691369

**Authors:** Hsuan Pai, Mauricio P. Contreras, Jose Salguero Linares, Daniel Lüdke, Andres Posbeyikian, Jiorgos Kourelis, Sophien Kamoun, Clemence Marchal

## Abstract

Following the perception of pathogen virulence proteins in plants, nucleotide-binding and leucine-rich repeat immune receptors (NLRs) are activated via a wide range of mechanisms. Singleton NLRs can both perceive effectors and trigger an immune response, whereas other NLRs specialise in either pathogen recognition (sensor NLRs) or activation of the immune response (helper NLRs). Sensor and helper NLRs can function as genetically linked pairs or in unlinked receptor networks. Although growing evidence suggests that NLRs conditionally oligomerise upon activation, our understanding of the resting state of NLRs prior to effector perception remains limited. Here, we investigated the oligomeric state of the genetically linked rice (*Oryza sativa*) sensor Pik-1 and helper Pik-2 NLR pair prior to effector activation when transiently expressed in *Nicotiana benthamiana* leaves. We show that both wild-type Pikm-1 and engineered Pikm-1^Enhancer^ sensors associate with Pikm-2 and form ∼1 MDa hetero-complexes in the resting state that accumulate at the plasma membrane. Our findings contribute to the growing evidence that pre-activation mechanisms vary widely across NLRs. This knowledge could be leveraged for disease resistance engineering strategies complementary to approaches focussing solely on effector binding.

**One sentence summary:** The Pik sensor-helper pair of rice immune receptor proteins forms a hetero-oligomer that accumulates at the plasma membrane in its resting state.

**Synopsis:** Nucleotide-binding and leucine-rich repeat receptors (NLRs) are key players in plant immune systems that recognise and respond to harmful pathogen effector proteins. NLRs can play specialised roles in either detecting effectors (sensor NLRs) or triggering the immune response (helper NLRs) and might function together as pairs or within larger networks. Although many NLRs are known to oligomerise upon the perception of effectors, much less is understood about their resting state prior to pathogen recognition. In this study, we investigated the resting state of a pair of rice NLRs, Pik-1 and Pik-2, transiently expressed in *Nicotiana benthamiana* leaves. These two proteins formed a sensor/helper complex at the cell membrane even in the absence of the effector protein. This discovery is significant because it reveals a new aspect of how these immune proteins are organised before activation, contributing to our understanding of the diverse ways plants prepare for immune responses. This finding adds to the growing body of evidence that, although many NLR proteins form complexes to respond to pathogens, their resting states and pre-activation mechanisms can vary widely.

## Introduction

Nucleotide-binding and leucine-rich repeat receptors (NLRs) are intracellular immune receptors found across all kingdoms of life^1,2^. In plants, NLRs directly or indirectly detect virulence proteins secreted by pathogens, termed effectors, which leads to the activation of an immune response and disease resistance^3^. Throughout evolution, NLR proteins have diversified from single functional units (functional singletons) capable of sensing pathogen effectors and initiating an immune response into specialised, sub-functionalized units comprising sensor NLRs that sense effectors or helper NLRs that mediate the downstream immune responses^4^. Across plant genomes, sensor and helper NLR genes can be genetically linked and cooperate as pairs, often in head-to-head orientation with a shared promoter sequence^4–12^. Additionally, some sensor and helper NLRs can function as complex receptor networks that are genetically unlinked^13,14^.

Plant NLRs exhibit a conserved domain architecture consisting of a variable N-terminal domain, a central nucleotide-binding domain shared by APAF1, certain *R* gene products, and CED-4 (NB-ARC) domain, and a C-terminal leucine-rich repeat region (LRR). Although the majority of NLRs have maintained this canonical domain architecture, they sometimes exhibit diverse modes of activation and signalling. An emerging paradigm in NLR biology is that NLRs conditionally oligomerise into inflammasome-like structures known as resistosomes upon activation^15^. However, our understanding of the resting and autoinhibited states of NLR proteins in plants prior to effector perception is limited. Moreover, much of our knowledge of plant NLRs is derived from the activation mechanisms of singleton and helper NLRs. How paired sensor and helper NLRs mechanistically function remains poorly understood. In this study, we addressed this question using the paired rice (*Oryza sativa*) immune proteins Pik-1 and Pik-2, which confer resistance to the rice blast pathogen *Magnaporthe oryzae* (syn. *Pyricularia oryzae*) and constitute a well-established model system for NLR receptor bioengineering^9,16–21^

NLRs can be phylogenetically classified based on their NB-ARC domains and the nature of their N-terminal domains. In flowering plants (angiosperms), these include Toll/Interleukin-1 receptor (TIR), G10 subclade coiled coil (CC_G10_), RESISTANCE TO POWDERY MILDEW 8 (RPW8) coiled coil (CC_R_), and coiled coil (CC) domains^22^. Some NLRs carry unconventional integrated domains (IDs) that are generally thought to mediate the detection of pathogen effectors either by directly binding to the effectors or by acting as a substrate for their enzymatic activity^5,6,23–26^. Several characterised sensor NLRs featuring an ID (NLR-IDs) function as a genetically linked pair, with a helper NLR that lacks an ID for immune activation upon effector detection^5,7,9,11,12^.

Despite the diversity observed across plant NLRs, a common theme among different classes of NLRs is the formation of higher-order oligomers termed resistosomes upon pathogen-induced activation. These resistosomes exhibit variable stoichiometry, with tetramers, pentamers, hexamers, and octamers described thus far^27–36^. The CC-NLR singletons ZAR1^27,28^ and Sr35^29,30^ form pentameric resistosomes at the plasma membrane, while CC_G10_-NLR WAI3 forms an octameric resistosome. By contrast, in asterid plants, the NRC (NLR required for cell death) network is composed of functionally specialized sensor and helper NLRs^13^, in which activation of sensor NLRs upon detection of pathogen effectors triggers the oligomerisation of helper NLRs and ultimately hypersensitive cell death and disease resistance^37–39^. Notably, NRC-dependent sensors are not part of the activated NRC oligomers and are thought to mediate oligomerisation of the helper NLR through an activate-and-release model via their central nucleotide-binding domain^40^. NRC2, NRC3 and NRC4 form hexameric resistosomes that accumulate at the plasma membrane^31,32,41^. Notably, NRC4, ZAR1, Sr35, and WAI3 display Ca^2+^ channel activity, which contributes to immune activation and cell death^29,31,34,42,43^. In another activation mechanism, TIR-NLRs act as sensors and oligomerise to form tetrameric holo-enzymes that produce nucleotide-derived second messengers upon effector detection. The newly generated small molecules induce heterodimerization of the lipase-like proteins enhanced disease susceptibility 1 (EDS1) with either senescence-associated gene 101 (SAG101) or phytoalexin deficient 4 (PAD4). Formation of EDS1-SAG101 or EDS1-PAD4 hetero-complexes is perceived by the CC_R_-NLR N REQUIREMENT GENE 1 (NRG1) or ACTIVATED DISEASE RESISTANCE 1 (ADR1) helper NLRs respectively, triggering their oligomerization^14,35^. Like singleton and NRC helpers, NRG1 and ADR1 oligomers show Ca^2+^-permeable cation channel activity at the plasma membrane^43^. Altogether, these findings indicate that the modes of activation of NLR proteins that lead to the formation of higher-order complexes are highly variable.

The diversity of NLR activation mechanisms is also reflected in their resting states. It is becoming increasingly apparent that NLR proteins adopt different conformations to remain in an (auto-)inhibited state and prevent autoactivation. For example, ZAR1 adopts a monomeric conformation in complex with its partner pseudokinase RKS1^27^ in its autoinhibited resting state. Interfering with ZAR1-RKS1 resting-state association does not trigger autoimmunity, suggesting that ZAR1 monomers are the minimal autoinhibited unit. Strikingly, the helper NRC2 forms homodimers^33^ and dimers of dimers (tetramers) that can assemble to form filament-like structures in its resting state^44^. Furthermore, recent blue native–polyacrylamide gel electrophoresis (BN-PAGE) assays indicated that the TIR-NLR pairs CHS3-CSA1 and RPS4-RRS1 appear to form heterodimers in their resting state, although the structure of the complexes remains to be elucidated^46,47^. Altogether, these observations suggest that oligomerisation is not exclusive to NLR activation and that oligomer formation in the resting state may constitute an additional layer of self-inhibition. Importantly, the diversity observed among the relatively few NLR resting states characterised so far suggests that many more resting-state configurations remain to be identified.

*Pikm-1* and *Pikm-2* are alleles of two genes at the *Pik* locus that encode a CC-NLR pair. This NLR pair confers resistance to strains of the rice blast fungus *M. oryzae* by detecting alleles of the AVR-Pik effector, including AVR-PikD^9,48–50^ (Figure 1A). The sensor Pikm-1 contains a non-canonical heavy-metal associated (HMA) integrated domain between its CC and NB-ARC domains that directly binds to AVR-Pik effectors. The HMA domain is the most variable domain across different Pik-1 alleles, suggesting that this domain is under selection imposed by AVR-Pik^48,51,52^. Given that the effector recognition specificity appears to be largely encoded by the integrated HMA in Pik-1, this integrated domain has been successfully targeted for immune receptor bioengineering to broaden or switch its effector recognition spectrum^19,20,53^. We recently demonstrated that replacing the Pikm-1 HMA domain with a completely unrelated sequence such as that of nanobodies that bind fluorescent proteins (FPs) resulted in a fully functional immune receptor able to sense FPs and trigger an effective immune response when co-expressed with the Pikm-2 helper^21^ (Figure 1A). Additionally, Pik-1 and Pik-2 can be successfully transferred from monocot to dicot species without requiring any additional components for the induction of an immune response^21^. Thus, the Pik CC-NLR pair is a highly promising NLR system for disease resistance engineering strategies across multiple crop species.

**Figure 1.**
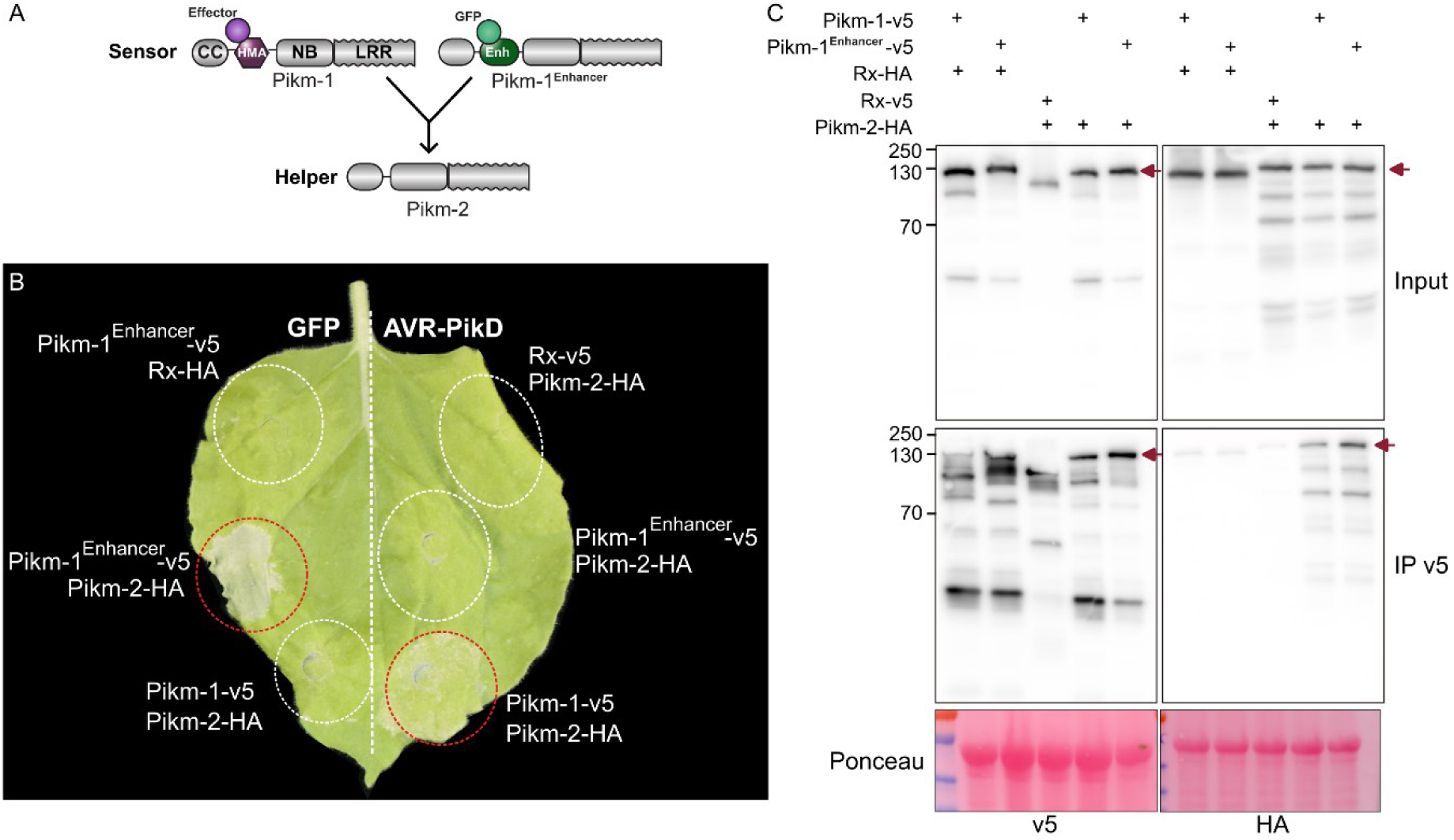
Pikm-1^Enhancer^ associates with Pikm-2 prior to effector activation. A) Diagram of the domain organisation of the sensors Pikm-1 and Pikm-1^Enhancer^ and the helper Pikm-2. Swapping the heavy-metal-associated (HMA) domain of Pikm-1 for the nanobody Enhancer, which binds to GFP^71^, leads to a functional receptor able to trigger an immune response in the presence of GFP^21^. Both the Pikm-1 and Pikm-1^Enhancer^ sensors signal through the Pikm-2 helper upon effector or ligand recognition to trigger an immune response. B) Pikm-2-6xHA, Pikm-1-2xV5, and Pikm-1^Enhancer^-2xV5 are functional in the presence of their ligand when transiently expressed in *N. benthamiana*. The image shows a representative leaf that was infiltrated with the indicated constructs and photographed 5 days after infiltration. Dashed circles indicate the infiltrated areas, and red circles show where a hypersensitive response was observed. C) Pikm-2 co-immunoprecipitates with both Pikm-1 and Pikm-1^Enhancer^ in the absence of their ligand. Rx-v5 + Pikm-2-HA and Rx-HA + Pikm-1-v5 were used as negative controls, and Pikm-1 + Pikm-2 was used as a positive control^16^. Total protein extracts (input) and proteins immunoprecipitated using v5 beads (IP v5) were immunoblotted with antibodies labelled at the bottom. Approximate molecular weights (kDa) are shown at left, and red arrows indicate bands corresponding to the Pik receptors. Ponceau staining of the membrane is shown below the immunoblots.

In a previous study, Zdrzałek et al.^16^ used co-immunoprecipitation assays to show that Pikm-1 and Pikm-2 associate *in planta* before and after effector-mediated activation^16^. However, the oligomeric state of such complexes has not been investigated. Determining the oligomeric state of the Pik complexes will deepen our understanding of how cooperating receptors trigger an immune response. Importantly, this would allow fine tuning of this receptor pair for disease resistance and complement current approaches focussing solely on effector binding. Notably, BN-PAGE assays have only recently been used to study plant NLR biochemistry^37,38,54,55^. Here, we used BN-PAGE to show that Pikm-1 and Pikm-2 form hetero-oligomeric complexes prior to effector-mediated activation in *Nicotiana benthamiana* leaves. We show that both wild-type Pikm-1 and Pikm-1^Enhancer 21^, a nanobody engineered variant that senses green fluorescent protein (GFP), form hetero-oligomers with Pikm-2 in their resting state. Taking advantage of mutations in the N-terminal domain of helper NLRs that prevent the induction of cell death upon effector activation^56^, we demonstrate that GFP, the ligand of Pikm-1^Enhancer^, is also part of the Pikm-1^Enhancer^-Pikm-2 hetero-oligomer. Additionally, when both proteins were expressed individually, we detected the helper Pikm-2 in both the membrane and soluble fractions in a membrane fractionation assay, and Pikm-1 mainly in the soluble fraction. Co-expression of the sensor and helper led to a shift in the localisation of Pikm-1 to the membrane and increased the membrane accumulation of Pikm-2. Our work emphasizes that NLR immune receptor proteins have diverse resting states and pre-activation mechanisms, despite their common feature of forming inflammasome-like resistosome oligomers during immune responses. Beyond developing a better understanding of immune receptor function, the details of these mechanisms lay the groundwork for developing new strategies in disease resistance engineering.

## Results

### Pikm-1^Enhancer^ associates with Pikm-2 prior to effector activation

We previously showed that Pikm-1^Enhancer^ specifically senses GFP and triggers an immune response in the presence of Pikm-2^21^. However, it is not known whether Pikm-1^Enhancer^ and Pikm-2 interact in the absence of their ligand in the same manner as Pikm-1 and Pikm-2^16^. To investigate this question, we transiently co-expressed Pikm-1^Enhancer^ and Pikm-2 in *N. benthamiana* leaves through agroinfiltration and performed co-immunoprecipitation (Co-IP) experiments (Figure 1B-C). Tagged Pikm-2 co-expressed with either Pikm-1 or Pikm-1^Enhancer^ triggered a hypersensitive response in the presence of the corresponding ligand (AVR-PikD for Pikm-1 and GFP for Pikm-1^Enhancer^, Figure 1B), demonstrating that these receptors are functional and not autoactive in the absence of their ligand under the tested conditions. Pikm-1^Enhancer^ co-immunoprecipitated with Pikm-2 to levels comparable to Pikm-1. By contrast, the negative control Rx (CC-NLR) co-immunoprecipitated only weakly with Pikm-1, Pikm-1^Enhancer^, or Pikm-2 (Figure 1C and Figure S1), corresponding to the background signal of the IP. We thus confirmed that Pikm-1^Enhancer^ and Pikm-2 follow the same interaction pattern as Pikm-1 and Pikm-2 in their resting state.

### The Pik pair forms a ∼1 MDa hetero-oligomer prior to activation

Although we observed an interaction between Pikm-1^Enhancer^-Pikm-2 and Pikm-1-Pikm-2 in their resting state, the oligomerisation state of these complexes is unknown. To examine the oligomerisation state of the Pik pair prior to effector-mediated activation, we co-expressed the Pikm-1+Pikm-2 or Pikm-1^Enhancer^+Pikm-2 pairs in *N. benthamiana* leaves and performed BN-PAGE assays. We observed a high-molecular-weight band at ∼1 MDa only in lanes where Pikm-1 or Pikm-1^Enhancer^ were co-expressed with Pikm-2 (Figure 2A, red asterisks). This band migrated at a similar size for both the sensor and helper (V5 and HA antisera blots, respectively). Because Pikm-1 and Pikm-2 are similar in size (∼110 kDa), it is impossible to determine whether the bands correspond to separate Pik homo-oligomers (a sensor-only and a separate, helper-only oligomer) or to a hetero-oligomer that includes both subunits (Figure 2B). Each sensor and helper was co-expressed with Rx as a negative control (Figure 2A, Figure S2).

**Figure 2.**
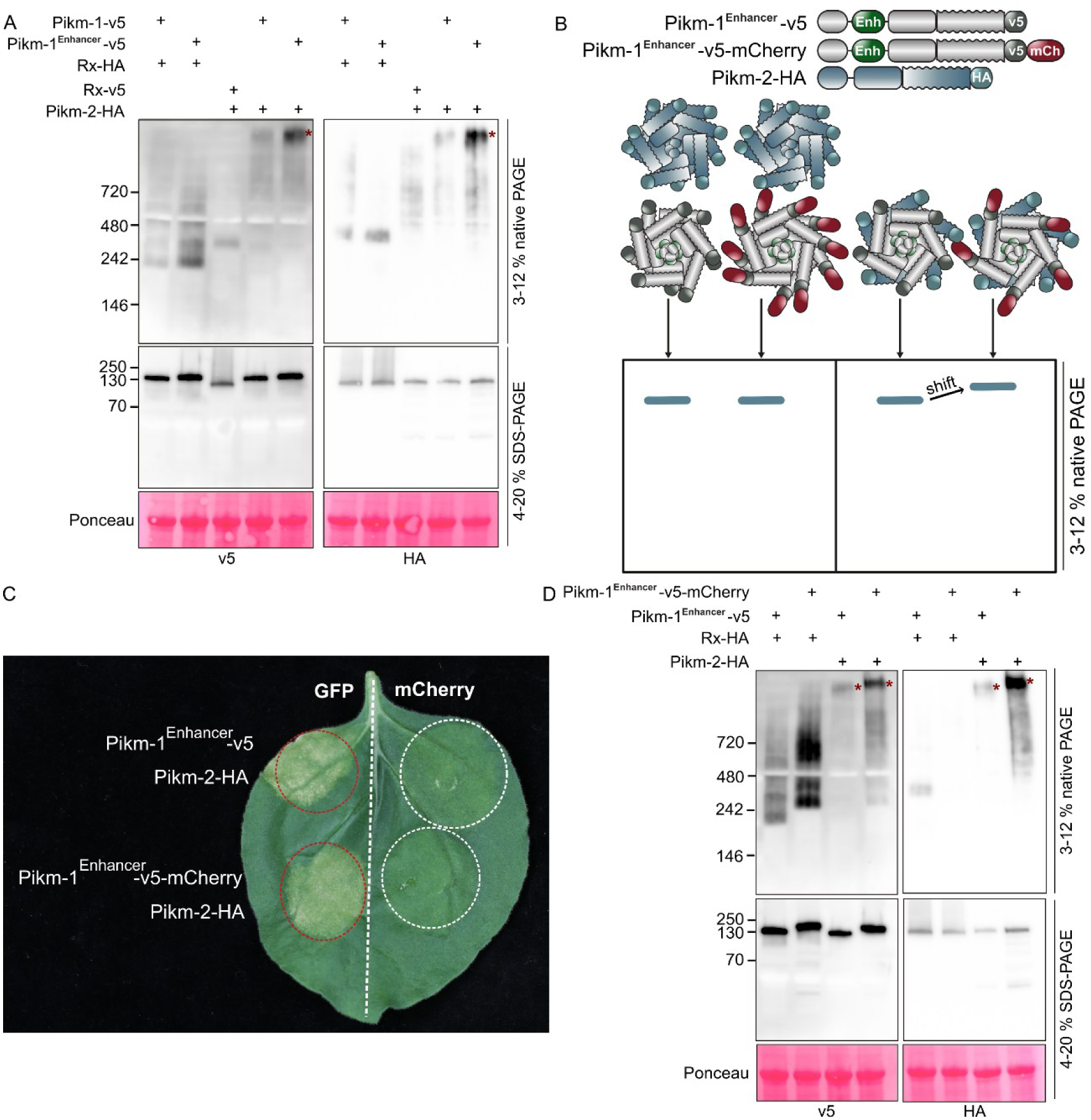
The Pik pair forms a ∼1 MDa hetero-oligomer prior to effector activation. A) BN-PAGE (top) and SDS-PAGE (middle) assays using *N. benthamiana* leaves co-expressing Pikm-2 with either Pikm-1 or Pikm-1^Enhancer^. Co-expression of Rx with Pikm-1 variants or Pikm-2 was used as a negative control. BN-PAGE and denaturing SDS-PAGE were run in parallel to verify the expression of all proteins. Approximate molecular weights (kDa) of the proteins are shown at left, and red asterisks indicate bands corresponding to the ∼1 MDa Pik oligomer. Bottom, Ponceau staining. B) Diagrams illustrating the strategy for using a ‘heavy’ Pikm-1^Enhancer^-2xV5-mCherry or ‘light’ Pikm-1^Enhancer^-2xV5 version of Pikm-1^Enhancer^. Briefly, a band shift of Pikm-2 in the immunoblot when using the ‘heavy’ Pikm-1^Enhancer^ variant indicates that both Pikm-2 and Pikm-1^Enhancer^ associate and are part of the same complex. C) Pikm-2-6xHA, Pikm-1-2xV5, Pikm-1^Enhancer^-2xV5, and Pikm-1^Enhancer^-2xV5-mCherry are functional when transiently co-expressed in the presence of their ligand in *N. benthamiana*. The image shows a representative leaf that was infiltrated with the indicated constructs and photographed 5 days after infiltration. Dashed circles indicate the infiltrated areas, and red circles show where a hypersensitive response was observed. D) BN-PAGE (top) and SDS-PAGE (middle) assays using ‘heavy’ Pikm-1^Enhancer^ and Pikm-2. Co-expression of Rx with Pikm-1 variants was used as a negative control. BN-PAGE and denaturing SDS-PAGE were run in parallel to ensure all components were expressed. Approximate molecular weights (kDa) of the proteins are shown on the left, and red asterisks point to the bands corresponding to ∼1 MDa Pik oligomer. Below, Ponceau staining showing Rubisco.

To resolve potential sensor and helper homo- or hetero-oligomers, we leveraged a previously published “heavy/light” tag approach^38^. We generated a C-terminally tagged tandem mCherry-V5 variant of Pikm-1^Enhancer^ termed “heavy”. This variant was functional and triggered a hypersensitive response in the presence of Pikm-2 and GFP, but not mCherry (Figure 2C). We tested whether this heavy variant of the Pikm-1^Enhancer^ sensor would induce a size shift of the Pikm-2 oligomer in BN-PAGE assays (Figure 2D). In the presence of Pikm-2, Pikm-1^Enhancer^ ‘heavy’ migrated as a higher-molecular-weight band in the BN-PAGE blot compared to C-terminally V5-tagged Pikm-1^Enhancer^ (termed ‘light’). Additionally, we observed an identical size shift in the BN-PAGE blot when probing for Pikm-2 (Figure 2D, red asterisks), suggesting that Pikm-1 and Pikm-2 are both part of the pre-activated ∼1 MDa oligomer.

We also observed intermediate bands at ∼242 and ∼480 kDa in the lanes where Pikm-1 or Pikm-1^Enhancer^ were expressed without Pikm-2 (Figure 2A and 2D). This suggests that the Pik sensors may form homo-oligomers in the absence of the corresponding Pik helper and that most of the Pikm-1 sensor pool shifts to the higher-order oligomeric state in the presence of the helper Pikm-2. Based on these results, we conclude that Pikm-1 and Pikm-2, as well as Pikm-1^Enhancer^ and Pikm-2, form a ∼1 MDa hetero-oligomer prior to activation that includes both the sensor and helper.

### Pikm-1^Enhancer^ and Pikm-2 preferentially accumulate in the membrane fraction when co-expressed

Growing evidence suggests that activated CC-NLRs and CC_R_-NLRs localise to membrane compartments^37,38,42,43,57,58^. Since the Pik sensor-helper pair forms a high-molecular-weight complex prior to activation, we investigated whether the Pik hetero-complex associates with membrane compartments in its resting state by performing membrane fractionation assays under non-denaturing conditions with the same constructs used for Co-IP and BN-PAGE assays. When Pikm-1^Enhancer^ was co-expressed with Rx, we detected Pikm-1^Enhancer^ in both the soluble and membrane fractions, with higher accumulation in the soluble fraction (Figure 3A, Figure S3). By contrast, Pikm-2 was present in comparable amounts in both the soluble and membrane fractions when co-expressed with Rx. Pikm-1^Enhancer^, but not Rx, accumulated in comparable amounts in both the soluble and membrane fractions upon co-expression with Pikm-2 (Figure 3A and S3, red asterisks). When co-expressed with Pikm-1^Enhancer^, Pikm-2 showed slightly stronger accumulation in the membrane fraction compared to co-expression with Rx (Figure 3A and S3, blue asterisks). We conclude that Pikm-1^Enhancer^ and Pikm-2 both accumulate in the soluble and membrane fractions, with preferential accumulation in the membrane fraction upon co-expression.

**Figure 3.**
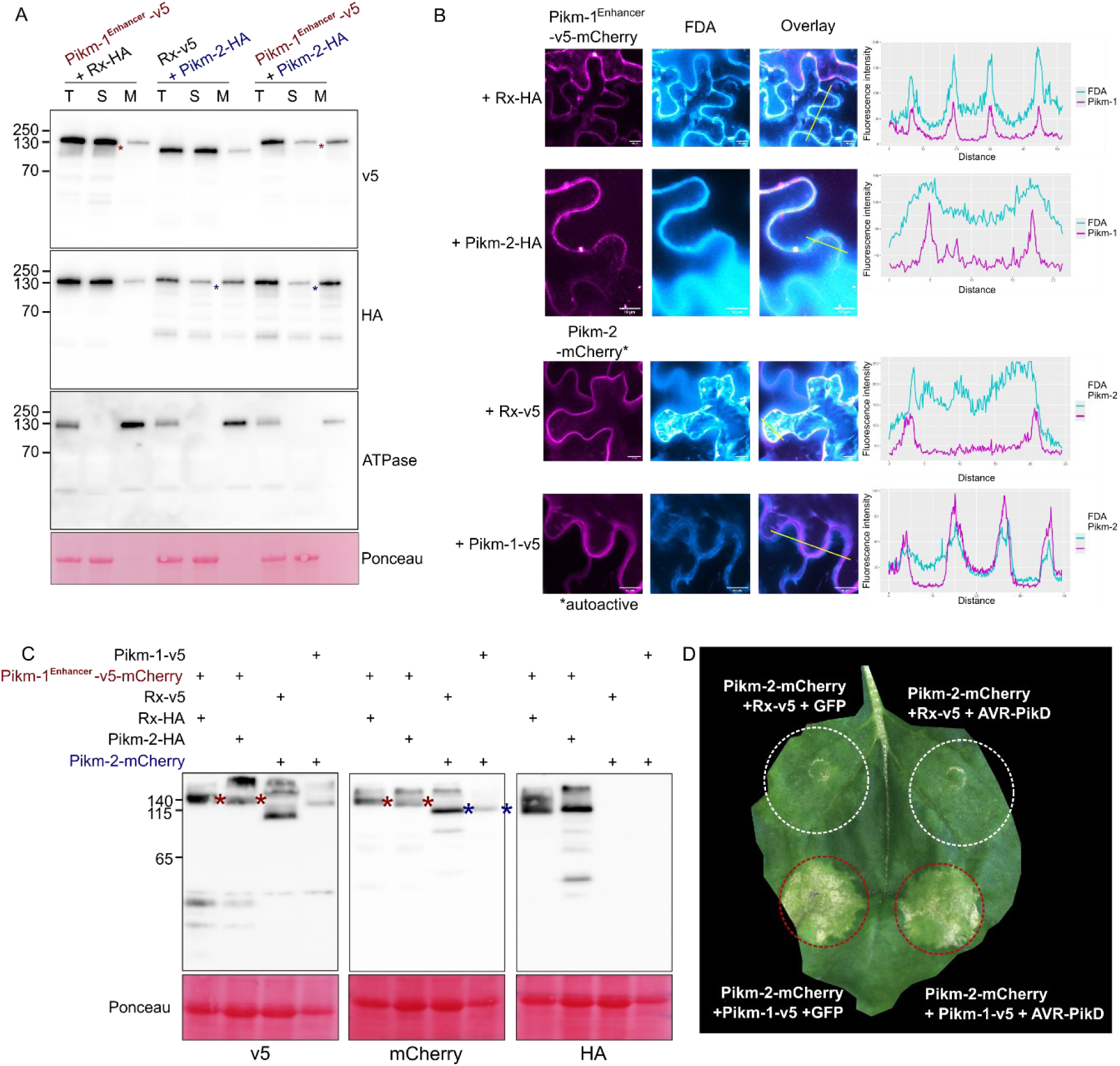
Pikm-1^Enhancer^ and Pikm-2 preferentially accumulate in the membrane fraction when co-expressed. A) Membrane fractionation assay of Pikm-1^Enhancer^ and Pikm-2. Rx was used as a negative control. Fractions: T = total, S = soluble, M = membrane. ATPase was used as a membrane marker. Rubisco was used as a marker for the total and soluble fractions and visualised by Ponceau staining. Red asterisks show the shift of Pikm-1^Enhancer^ from the soluble to the membrane fraction upon co-expression with Pikm-2. B) Confocal microscopy images of maximum projection of *z*-stacks of 10–25 slides showing the subcellular localisation of Pikm-1^Enhancer^-v5-mCherry or Pikm-2-mCherry co-expressed with Rx or the matching NLR from the Pik pair. Fluorescein was used as a cytosolic marker. The graphs at right depict the fluorescence intensity from each channel across the yellow line shown on the overlay image. All components were transiently expressed in *N. benthamiana* via agroinfiltration, and subcellular localisation was observed at 2 dpi. Laser excitation was set to 561 nm (diode) for mCherry and to 488 nm (argon) for fluorescein, and detection wavelengths were set to 590–660 nm and 500–560 nm for mCherry and fluorescein, respectively. C) SDS-PAGE assay of the samples used for confocal microscopy. Total protein extracts (input) were immunoblotted with the antibodies labelled at the bottom. Approximate molecular weight (kDa) of the protein is shown at left. Ponceau staining showing Rubisco is shown below the immunoblots. D) Cell death assay of *N. benthamiana* leaves co-expressing Pikm-2-mCherry with Pikm-1-v5 or Rx-v5 and GFP (negative control) or AVR-PikD, an effector from *M. oryzae* recognised by the Pikm pair. The image shows a representative leaf that was infiltrated with the indicated constructs and photographed 5 days after infiltration. Dashed circles indicate the infiltrated areas, and red circles show where a hypersensitive response was observed.

### Pikm-1^Enhancer^ and Pikm-2 localise to both the cytosol and plasma membrane when co-expressed

To investigate the subcellular localisation of Pikm-1^Enhancer^ and Pikm-2, we used confocal microscopy (Figure 3B). Since Pikm-1^Enhancer^ triggers cell death in the presence of GFP, CFP, YFP and mNeonGreen, we co-expressed Pikm-1^Enhancer^-v5-mCherry (Pikm-1^Enhancer^ ‘heavy’, Figure 2) with Pikm-2-HA or Rx-HA in *N. benthamiana* leaves to determine its subcellular localisation. For Pikm-2, we generated a C-terminally tagged version with mCherry to track its localisation upon co-expression with Pikm-1^Enhancer^-v5 or Rx-v5. Since Pikm-2-mCherry becomes autoactive at 3 dpi in the presence of Pikm-1^Enhancer^-v5 but not Rx-v5 (Figure 3D), we performed confocal imaging at 2 dpi before cell death was visible, the same time point used for the membrane fractionation assays (Figure 3A). We incubated leaf disks in a 0.5 µM fluorescein diacetate (FDA) solution for 10 minutes prior to imaging. FDA is membrane permeable and accumulates in viable cells as fluorescein, providing a green fluorescent marker for the cytosol.

Pikm-1^Enhancer^ fully co-localised with fluorescein when co-expressed with Rx, as indicated by the presence of cytosolic strands. When co-expressed with Pikm-2, Pikm-1^Enhancer^ remained cytoplasmic, but puncta were occasionally observed. In the absence of Pikm-1^Enhancer^, Pikm-2 co-localised with fluorescein at the cell periphery, with few or no cytosolic strands detected in maximum projection images derived from 10–15 slices. These observations indicate that Pikm-2 associated with the plasma membrane in the presence of both Rx and Pikm-1^Enhancer^ and was also present in the cytosol. By contrast, Pikm-1^Enhancer^ clearly localised to the cytosol when co-expressed with Rx and occasionally formed puncta in the presence of Pikm-2, suggesting that Pikm-2 can alter the subcellular localisation of Pikm-1^Enhancer^.

We then compared the localisation of Pikm-1 and Pikm-2 to that of the plasma membrane marker LTI6B from *Arabidopsis thaliana*^59^ (Figure S4). Because Pikm-1^Enhancer^ triggers cell death in the presence of free GFP, CFP, YFP, and mNeonGreen, we used wild-type Pikm-1 fused to mNeonGreen at its C-terminus for these assays. Pikm-2 was fused to mNeonGreen as well, while LTI6B was fused to mCherry, both at their C-termini. Co-expression of Pikm-1–mNeonGreen or Pikm-2–mNeonGreen with Rx resulted in autoactivity, which was not observed with the Pikm-1–mNeonGreen + Pikm-2–HA or Pikm-2–mNeonGreen + Pikm-1–v5 combinations. Therefore, we performed confocal imaging at 2 dpi, before cell death became apparent.

Consistent with observations for Pikm-1^Enhancer^, Pikm-1 co-localised with cytosolic strands and partially co-localised with the plasma membrane marker LTI6B at the cell periphery when co-expressed with Rx (Figure S4). When co-expressed with Pikm-2, Pikm-1 remained associated with cytosolic strands and co-localised with LTI6B at the cell periphery. Additionally, we detected Pikm-1 puncta in a small proportion of cells when Pikm-1 was co-expressed with Pikm-2, but not with Rx. Pikm-2 co-localised with LTI6B and formed puncta in a small proportion of cells when co-expressed with Pikm-1 or with Rx (Figure S4). Together, these results are consistent with the findings from membrane fractionation assays, supporting the conclusion that the Pikm-1^Enhancer^/Pikm-2 and Pikm-1/Pikm-2 hetero-complexes primarily localise to the plasma membrane, with a pool present in the cytosol.

### GFP associates with the Pikm-1^Enhancer^-Pikm-2 hetero-complex

Finally, we investigated whether the Pik hetero-complex would undergo any changes upon treatment with a ligand that binds to Pik-1. We could not perform these experiments using wild-type Pik proteins because hypersensitive cell death interferes with the biochemical assays. We therefore took advantage of the occurrence of the MADA motif in the N-terminal α1 helix of Pikm-2^56^ to introduce two mutations into this motif (Pikm-2^L19E,L23E^, subsequently termed Pikm-2^EE^) that abolished the induction of cell death upon co-expression with Pikm-1^Enhancer^ and GFP (Figure 4A). Mutations in this motif have previously been used to prevent the induction of cell death without compromising NLR activation and resistosome formation, enabling biochemical studies^38,56^.

**Figure 4.**
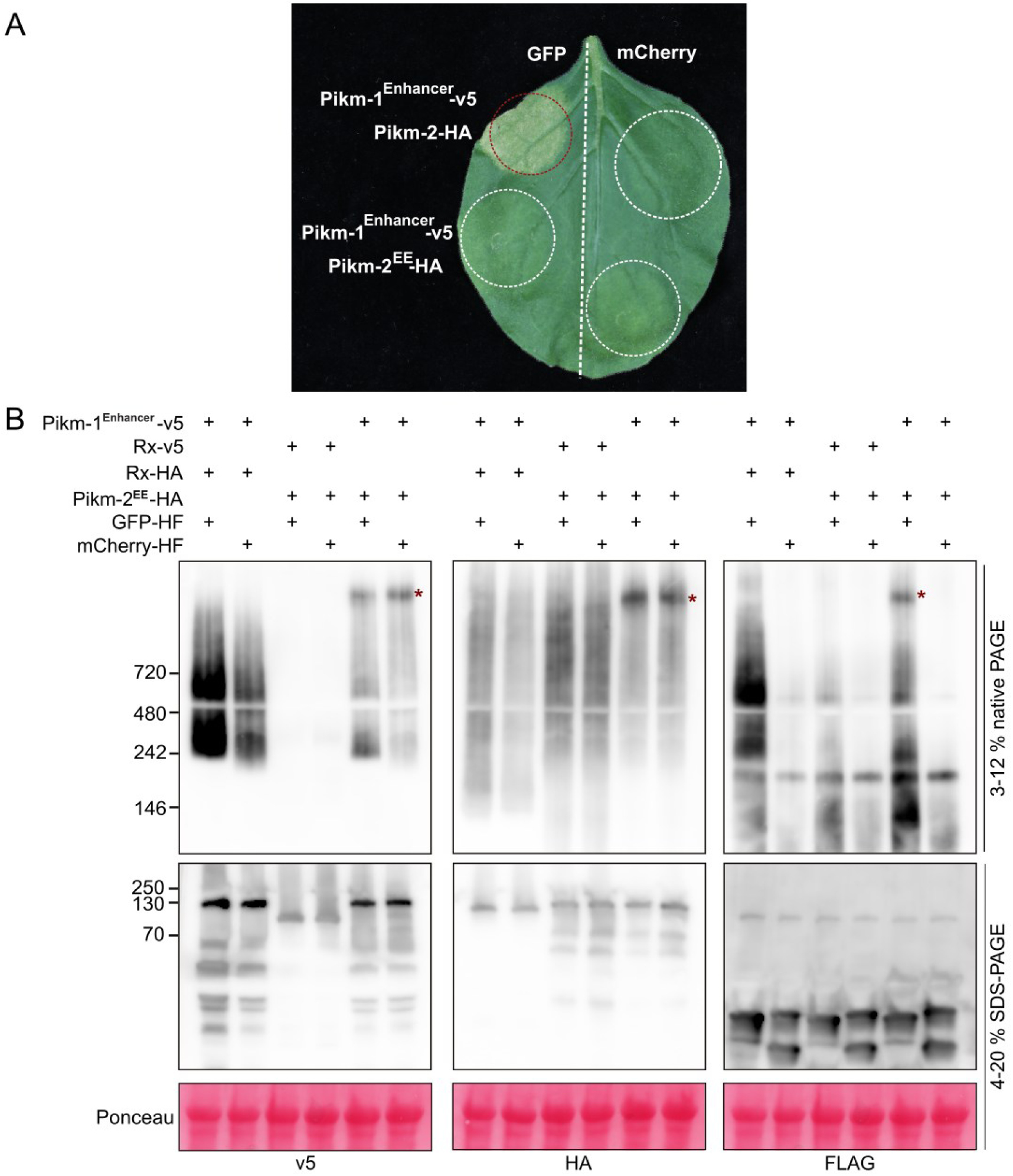
The GFP ligand associates with the Pikm-1^Enhancer^/Pikm-2 hetero-complex. A) Cell death assay of *N. benthamiana* leaves co-expressing Pikm-1 ^Enhancer^ with Pikm-2 or Pikm-2^EE^ with GFP (ligand) or mCherry (negative control). The image shows a representative leaf that was infiltrated with the indicated constructs and photographed 5 days after infiltration. Dashed circles indicate the infiltrated areas, and red circles show where a hypersensitive response was observed. B) BN-PAGE (top) and SDS-PAGE (middle) assays of Pikm-2^EE^ + Pikm-1^Enhancer^. Co-expression of Rx with Pikm-1 ^Enhancer^ or Pikm-2^EE^ was used as a negative control for hetero-oligomer formation. mCherry was used as a negative control for the presence of the ligand bound to Pikm-1^Enhancer^. BN-PAGE and SDS-PAGE samples were run in parallel to ensure all components were expressed. Approximate molecular weights (kDa) of the proteins are shown on the left, and red asterisks point to the bands corresponding to the ∼1 MDa Pik oligomer. Below, Ponceau staining showing Rubisco.

To determine whether treatment with the ligand would alter the Pik hetero-complex, we co-expressed Pikm-1^Enhancer^ and Pikm-2^EE^ with GFP or the negative control mCherry in *N. benthamiana* leaves and performed BN-PAGE assays (Figure 4B and Figure S5). We detected a GFP signal (using anti-FLAG antibodies) at ∼1 MDa only when GFP was co-expressed with Pikm-1^Enhancer^ and Pikm-2^EE^. By contrast, mCherry did not yield a signal at the ∼1 MDa Pikm-1^Enhancer^ /Pikm-2^EE^ complex. We conclude that GFP was integrated into the ∼1 MDa Pikm-1^Enhancer^/Pik-2 hetero-oligomer complex.

Despite these findings, we could not draw any conclusions about the activation state of the Pik NLR pair. Given that the ∼1 MDa bands were identical regardless of the presence of GFP, we could not conclude that the Pik pair was activated by GFP or whether Pikm-2^EE^ is a non-functional mutant of Pikm-2 that is blocked in a conformation similar to that of its resting state.

## Discussion

In this study, we investigated the oligomeric state and subcellular localisation of the CC-NLR pair Pikm-1 and Pikm-2 prior to activation. We leveraged a previously described bioengineered Pikm-1 variant, Pikm-1^Enhancer^, which specifically recognises GFP^21^ and enhances the accumulation of the Pikm-1/Pikm-2 complex in BN-PAGE assays compared with the wild type in our experimental conditions (Figure 2A). We demonstrated that Pikm-1 and Pikm-2, as well as Pikm-1^Enhancer^ and Pikm-2, form hetero-oligomers in their resting state (Figure 1 and 2). This hetero-oligomer accumulates at the plasma membrane (Figure 3). Furthermore, we generated the Pikm-2^EE^ mutant, which is unable to trigger cell death in the presence of Pikm-1^Enhancer^ and its ligand GFP, to determine whether the ligand is part of the hetero-oligomer. GFP migrated at the same molecular weight as the Pikm-1^Enhancer^-Pikm-2^EE^ hetero-oligomer, indicating that GFP may directly associate with the Pik hetero-complex (Figure 4). Overall, our study revealed that the sensor and helper of the CC-NLR Pik pair accumulate at the plasma membrane in the form of a ∼1 MDa hetero-complex in its resting state, highlighting the diversity of NLR activation mechanisms.

### The Pik pair forms a hetero-oligomer in its resting state

There is growing evidence that CC-NLRs remain autoinhibited in their resting state through a variety of mechanisms. For example, Arabidopsis ZAR1 exists as a monomer in complex with additional host components termed “guardees” prior to activation^27^. Additionally, we previously showed that NRC2 from *N. benthamiana* and tomato (*Solanum lycopersicum*) forms a homodimer in its resting state^33,44^. In its resting state, *Sl*NRC2 also forms higher-order dimer-based oligomers, leading to the formation of homo-tetramers and filaments, although these states are not required for NRC2 activation^44^. High-order oligomeric resting states were also observed for the mammalian NLR protein NLRP3 (NACHT-, leucine-rich-repeat-, and pyrin domain-containing protein 3)^60,61^. Prior to activation, NLRP3 forms oligomeric double-ring cages bound to membranes. Importantly, this conformation is required for the activation of NLRP3^60^. Here, we showed that the Pik sensor-helper pair forms a high-order hetero-oligomer in its resting state. Notably, hetero-oligomeric assembly was also reported for the TIR-NLR sensor pairs CSA1-CHS3 and RPS4-RRS1 in the resting state, which also remained in complex following effector activation^46,47^. Determining whether the Pikm-1-Pikm-2 oligomer changes its conformation or oligomeric state following ligand perception upon activation will require additional experiments, ideally using structural biology approaches. However, determining the structures of NLR pairs has been challenging, and their structures have not yet been resolved.

Although we demonstrated that both the Pikm-1 sensor and the Pikm-2 helper are part of the Pik sensor-helper pre-activation complex, the stoichiometry of the hetero-oligomer remains unclear. In BN-PAGE assays, we observed bands at ∼242 kDa and between 480 and 720 kDa when the sensors Pikm-1 or Pikm-1^Enhancer^ were expressed without Pikm-2 (Figure 2, Figure 4, Figure S2), indicating that the sensor alone may either form homo-oligomers of different stoichiometries or associates with unknown host components conserved between the heterologous *N. benthamiana* system and rice. By contrast, we did not consistently observe similar complexes for Pikm-2 in the absence of Pikm-1, although two potential Pikm-2 oligomeric states were visible in the BN-PAGE blots (Figure 2B, Figure S5). These observations raise the question of whether the final Pik resting state hetero-oligomer forms through the association of pre-formed sensor and helper homo-oligomers or through the oligomerisation of sensor-helper heterodimers. Furthermore, if the GFP-bound Pikm-1^Enhancer^-Pikm-2^EE^ hetero-oligomer reflects the activated state of the Pik pair, how does ligand binding trigger its activation and subsequent cell death? Resolving the structures of the Pik oligomer in both the resting and activated states will be essential to address these questions.

### The Pik hetero-oligomer accumulates at the plasma membrane

Using both membrane fractionation and confocal microscopy, we observed that Pikm-1 shifted from the cytosol to the plasma membrane when expressed with Pikm-2 (Figure 3, Figure S3 and S4). This indicates that Pikm-2 is the main driver for plasma membrane localization in the complex. This helper protein exhibits similar subcellular dynamics to *At*ADR1, *At*NRG1, and *At*RPM1, which also associate with the plasma membrane in their resting state^62,63^. How does Pikm-2 recruit Pikm-1 to the plasma membrane? Perhaps the two proteins use the plasma membrane as an oligomerisation scaffold, similar to what was observed for the cage-like structures that shield NLRP3 from mis-activation^60^. Further experiments are needed to determine the biological relevance of the plasma membrane association of the Pik complex in the resting state and whether it is required for the activation of the immune response upon effector recognition.

Several studies have shown that effector-activated or autoactive NLRs form puncta that associate with the plasma membrane^38,43,57,64^. These puncta are thought to be resistosomes or aggregates of membrane-associated resistosomes that actively contribute to the onset of cell death upon activation. We observed that both Pik-1 and Pik-2 occasionally formed puncta in *N. benthamiana* cells (Figure 3B, Figure S4). Specifically, only autoactive Pikm-2-mNeonGreen formed puncta in the absence of Pikm-1, while Pikm-2-mCherry did not. Pikm-1 and Pikm-1^Enhancer^ only formed puncta when co-expressed with Pikm-2 (Figure 3B, Figure S4). Given that we only observed puncta in a small fraction of the cells expressing these components and that most of the NLR-fluorescent fusion proteins used in these experiments were autoactive, we cannot draw conclusions about the biological relevance of the Pik puncta at this time. Overall, the biochemical nature of these puncta remains elusive, as they might consist of resistosome foci or clusters of inactive NLR complexes.

### Outlook

The Pik NLR pair is present as an allelic series in rice cultivars and confers resistance against *M. oryzae*, the causal agent of rice blast disease, by recognising specific alleles of the AVR-Pik effector^9,48–50^. Recent studies have shown that the *Pik-1* and *Pik-2* alleles have co-evolved to tightly control the immune response^20,51^. For instance, mismatching the allelic sensor Pikp-1 with the helper Pikm-2 caused hypersensitive cell death in the absence of the effector, while co-expressing Pikm-1 with Pikp-2 resulted in a weakened immune response^51^. Compatibility between Pik alleles appears to be driven by specific polymorphic residues, such as Glu230 located in the NB-ARC domain of Pik-2^51^, and variations within the integrated HMA domain of Pik-1^20^. Notably, allelic mismatching did not prevent hetero-complex formation in Co-IP experiments, although co-evolved pairs showed a preferential association^51^. This raises several important questions: Does the binding affinity of these sensor-helper alleles directly affect their hetero-oligomeric state? Which molecular determinants underlie constitutive cell death or reduced activity in mismatched Pik hetero-oligomers? Can incompatibility at the sensor-helper hetero-oligomerisation interface lead to autoactivation of the Pik system? Answering these questions will provide insights into how Pik-1 and Pik-2 have co-adapted and how co-evolution influences the oligomerisation abilities of different alleles. Furthermore, a better understanding of the molecular basis of sensor-helper compatibility in this system may guide future Pik bioengineering efforts.

The resting state of the Pik hetero-oligomer adds to the diversity of CC-NLR activation mechanisms characterised to date. It will be interesting to determine whether other genetically linked CC-NLR pairs also form hetero-oligomers in the resting state. It is likely that additional diversity in CC-NLR pairs in the resting state remains to be discovered, especially since not all characterised CC-NLR pairs function cooperatively, as does the Pik system^16^. For instance, unlike Pik-1 and Pik-2, the rice Pia pair (RGA4 and RGA5) appears to function via negative regulation, with the sensor RGA5 maintaining its helper RGA4 in a repressed state. Effector binding by RGA5 is thought to trigger a conformational change in RGA5 and subsequent release of RGA4 inhibition leading to the induction of cell death in *N. benthamiana*^65^. Although RGA4 and RGA5 interact in both the absence and presence of the effector AVR-Pia, whether this negative regulation model influences the resting state or activated oligomerisation profile of this NLR pair remains to be determined.

We propose a model describing the cooperation between a sensor and a helper in a CC-NLR pair via the formation of a large (∼1 MDa) membrane-associated hetero-oligomer in the resting state (Figure 5). This work adds to the growing evidence that, while oligomerisation is a hallmark of NLR-mediated immune responses, the resting states and mechanisms leading to NLR assembly into resistosomes exhibit remarkable diversity. These findings should advance the field beyond a uniform view of NLR structure and function and further facilitate comparative analyses of NLRs across diverse plants.

**Figure 5.**
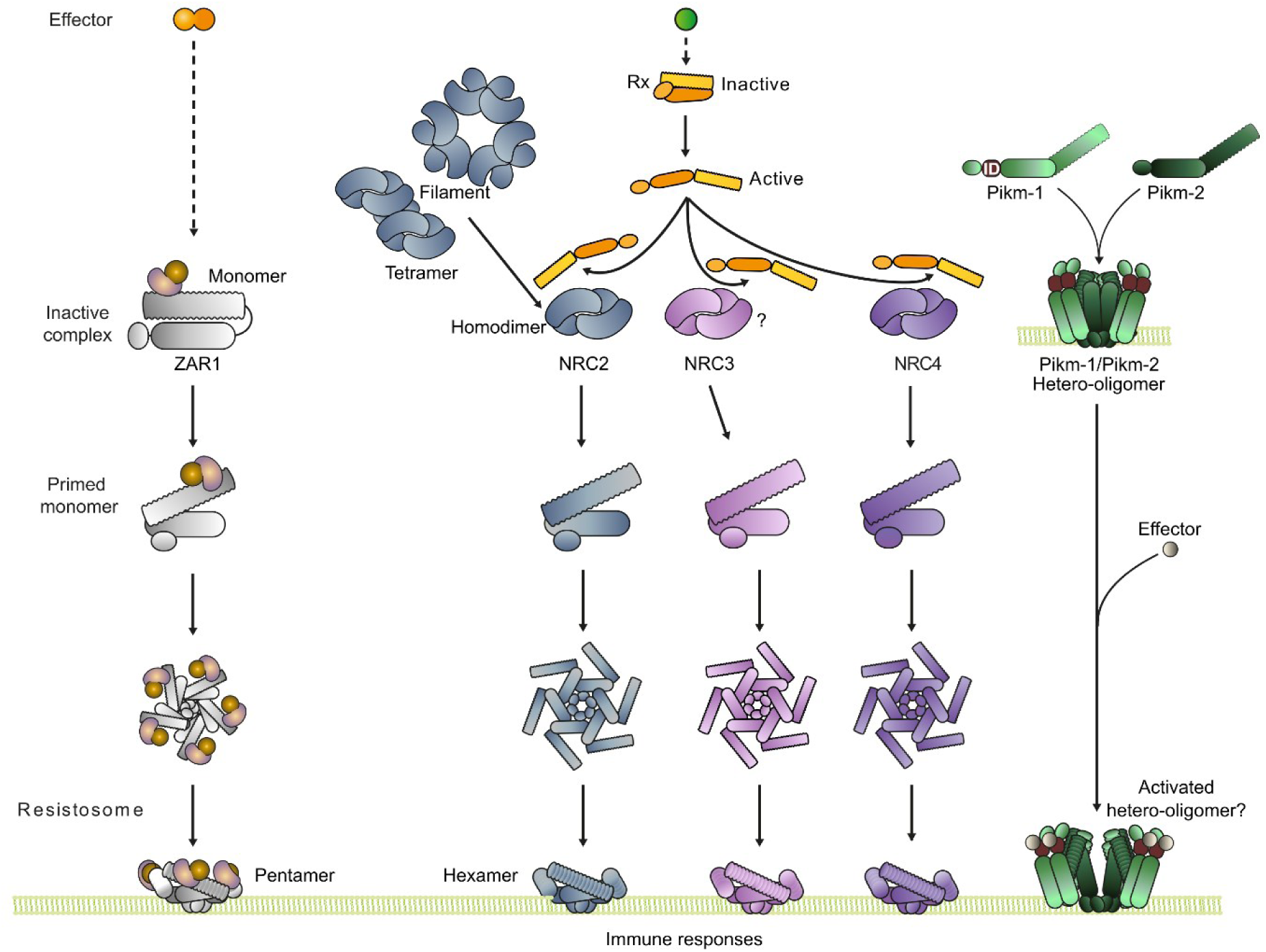
CC-NLRs exhibit diverse conformations in the resting state. Updated figure from Selvaraj et al.^72^ depicting different NLR resting and activated states. While ZAR1 exists as a monomer bound to its guardees in its resting state, several forms have been observed for the NRC2 helper^44,72^, including dimers, tetramers, and filaments. While filament-like structures were observed via confocal microscopy for NRC2 upon overexpression in planta, these structures were not detected for its paralog NRC4^72^, suggesting a potentially different resting state configuration for NRC4. Our work suggests that the Pik CC-NLR pair forms a hetero-complex bound to the plasma membrane in its resting state. How effector binding to the integrated domain (ID) of the Pik-1 sensor leads to the activation of the complex remains to be determined.

## Material and Methods

### Plant growth conditions

Wild-type *N. benthamiana* plants were grown in a controlled environment growth chamber with a temperature range of 22–25°C, relative humidity of 45–65%, and a 16/8-h light/dark cycle.

### Plasmid construction

The Golden Gate Modular Cloning (MoClo) kit^66^ and the MoClo Plant Parts kit^67^ were used for cloning unless otherwise specified (Table S1). Cloning design and sequence analysis were performed using Geneious Prime (v2022.2.2; https://www.geneious.com). For the mRFP-Lti6b-TurboID plasma membrane marker, the coding sequence of *LOW-TEMPERATURE-INDUCED6b* from Arabidopsis (*Lti6b*, GenBank accession number AT3G05890) was amplified without the stop codon alongside the TurboID sequence from AT3G05890. Both PCR fragments were ligated using overlapping PCRs. The fusion construct *Lti6b*-TurboID was then cloned into the pENTR/D-TOPO plasmid (Thermo Scientific). The final RFP-Lti6b-TurboID construct was generated via Gateway cloning (LR reaction, Thermo Scientific) of pENTR/D-TOPO into pGWB555^68^. Plasmid construction is described in Table S1.

### Cell death assays in *N. benthamiana*

Transient gene expression in *N. benthamiana* was performed by agroinfiltration as described by van der Hoorn et al. ^69^. Briefly, *Agrobacterium tumefaciens* strain GV3101 pMP90 carrying binary vectors was inoculated from glycerol stock in LB medium supplemented with the appropriate antibiotics and grown overnight at 28°C until saturation. The cells were harvested by centrifugation at 2000 × *g* at room temperature for 5 min. The cells were washed once and resuspended in infiltration buffer (10 mM MgCl_2_, 10 mM MES-KOH pH 5.6, 200 μM acetosyringone) to the appropriate OD_600_ (see Table S1) in various combinations and incubated for 30 min at room temperature prior to infiltration into 4- to 5-week-old *N. benthamiana* leaves. Two leaves from three plants were inoculated per experiment (*N* = 6), and the experiment was repeated three times. In all assays where the original Pikm was co-infiltrated with its cognate effector AVR-PikD, the Tomato bushy stunt virus (TBSV) silencing inhibitor p19^70^ was co-infiltrated into the leaves to enhance expression and ensure a reliable hypersensitive response. Hypersensitive cell death phenotypes were visually scored 4–5 days post infiltration in a range from 0 (no visible necrosis) to 7 (fully confluent necrosis) according to Adachi et al.^56^.

### Protein extraction for BN- and SDS-PAGE assays

Six *N. benthamiana* leaf discs (9 mm diameter) collected at 2 days post agroinfiltration were homogenised in 200 μL extraction buffer (10% glycerol, 50 mM Tris-HCl [pH 7.5], 5 mM MgCl_2_, 50 mM NaCl) supplemented with 10 mM DTT, 1× protease inhibitor cocktail (SIGMA), and 0.5% Triton-X (SIGMA) and incubated on ice for 10 min. The samples were centrifuged at 5,000 × *g* for 15 min at 4°C, and the supernatant was used for BN- and SDS-PAGE assays.

### BN-PAGE assays

For BN-PAGE, 25 μL of the supernatant obtained from protein extraction described above was diluted as per the manufacturer’s instructions by adding NativePAGE 5% G-250 sample additive, 4× Sample Buffer, and water. After dilution, the samples were run on Native PAGE 3–12% Bis-Tris gels alongside a Native Marker (SERVA). The separated proteins were transferred to polyvinylidene difluoride membranes using NuPAGE Transfer Buffer with a Trans-Blot Turbo Transfer System (Bio-Rad) as per the manufacturer’s instructions. Proteins were fixed to the membranes by incubating in 8% acetic acid for 15 min, washed with water, and left to dry. The membranes were subsequently re-activated with ethanol to correctly visualise the unstained native protein marker. The membranes were immunoblotted as described below.

### SDS-PAGE assays

For SDS-PAGE, 25 µL of the supernatant obtained from protein extraction described above was diluted in 75 µL 4× SDS-PAGE sample buffer (final concentration: 50 mM Tris-HCl [pH 6.8], 100 mM DTT, 2% SDS, 0.01% bromophenol blue, 10% glycerol) and incubated at 72°C for 10 min. The samples were centrifuged at 5,600 × *g* for 2 min, and 10 µL of the supernatant was separated on a 10–20% SDS-PAGE gel (Bio-Rad) and transferred onto a polyvinylidene difluoride (PVDF) membrane using a Trans-Blot turbo transfer system (Bio-Rad). The membranes were immunoblotted as described below.

### Co-immunoprecipitation

Two leaves from the bottom half of a *N. benthamiana* plant (∼1 g) collected 2 days post agroinfiltration were homogenised in 2:1 (mL:g) buffer:sample extraction buffer (10% glycerol, 25 mM Tris-HCl, pH 7.5, 1 mM EDTA, 150 mM NaCl, 2% [w/v] PVPP 10 mM DTT, 1× protease inhibitor cocktail [SIGMA], 0.2% IGEPAL® CA-630 [SIGMA]). The samples were incubated on ice for 10 min and centrifuged at 5,000 × *g* for 30 min. The supernatant was passed through a Minisart 0.45 μM filter (Sartorius Stedim Biotech, Goettingen, Germany), and 25 µL of the sample was used for SDS-PAGE as described above. For immunoprecipitation, 1 mL of filtered supernatant was mixed with 60 μL v5 beads (Roche) that had been washed three times and resuspended in immunoprecipitation buffer at a 1:1 ratio (10% glycerol, 25 mM Tris-HCl, pH 7.5, 1 mM EDTA, 150 mM NaCl, 0.2% IGEPAL® CA-630 [SIGMA]). The beads were washed six times with immunoprecipitation buffer and resuspended in 50 μL 4× SDS-PAGE sample buffer. The samples were incubated at 72°C for 10 min to release the proteins from the beads and centrifuged at 5,600 × *g* for 2 min to pellet the beads. Finally, 15 μL of supernatant was loaded on a 10–20% SDS-PAGE gel (Bio-Rad) and transferred onto a PVDF membrane using a Trans-Blot turbo transfer system (Bio-Rad). The membranes were immunoblotted as described below.

### Confocal microscopy

Transient gene expression in *N. benthamiana* was performed by agroinfiltration as described above. Leaf disks (0.4 mm diameter) were directly mounted in 0.5 μM Fluorescein diacetate (FDA) solution and incubated for 10 min in the dark before imaging for the experiments shown in Figure 3. For Figure S4, LTI6B from Arabidopsis fused to mRFP at its N-terminus was used as a plasma membrane marker. The abaxial epidermal tissue was imaged under a Zeiss LSM880 confocal microscope using a 40× water immersion objective lens. Laser excitation was set to 561 nm (diode) for mCherry and to 488 nm (Argon) for fluorescein (Figure 3) or mNeonGreen (Figure S5). *Z*-stacks were processed in Zen lite (v3.12) to generate maximum projections and the fluorescence intensity profiles shown in Figure 3 and Figure S5. Images shown are representative of three experiments in which two plants per condition were infiltrated, and leaf disk were sampled from two leaves per plant.

### Membrane fractionation assays

Membrane fractionation assays were performed as previously described^38^. Briefly, leaf material was ground into a fine powder in liquid nitrogen and combined with 2 volumes of extraction buffer (0.81 M sucrose, 5% [v/v] glycerol, 10 mM EDTA, 10 mM EGTA, 5 mM KCl, and 100 mM Tris-HCl [pH 7.5]) supplemented with 5 mM DTT, 1% Sigma Plant Protease Inhibitor Cocktail, 1 mM PMSF, and 0.5% PVPP. The samples were vortexed for 1 min, and cell debris was pelleted via two subsequent centrifugation steps at 1,000 × *g* for 5 min. The supernatant was diluted 1:1 in distilled water, and an aliquot of the diluted sample was separated as the total fraction (T). The remaining supernatant (200–300 μl) was centrifuged at 21,000 × *g* for 90 min at 4°C. This centrifugation step separated the sample into the soluble fraction (S) in the supernatant and the membrane-enriched fraction (M) in the pellet. After the soluble fraction was set aside, the pellet was resuspended in diluted extraction buffer without PVPP. The fractions were diluted with SDS loading dye, and proteins were denatured by incubating at 50°C for 15 min. Immunoblotting was performed as described in the SDS-PAGE section. Endogenous plasma membrane ATPase was detected using anti-H + ATPase (AS07 260) antibody (Agrisera) as a control to confirm successful membrane enrichment.

### Immunoblotting

Membranes were blocked in 3% dried milk dissolved in Tris-buffered saline (50 mM Tris-HCl [pH7.5], 150 mM NaCl) supplemented with 1% Tween® for 20 to 30 min. The membranes were probed with either rat monoclonal anti-HA antibody (3F10, Roche; 1:5000 dilution), mouse monoclonal ANTI-FLAG® antibody conjugated to HRP (M2, Sigma; 1:5000 dilution), or mouse monoclonal anti-V5 antibody (V5-10, Sigma-Aldrich; 1:5000). After adding either Pierce ECL Western Substrate (Thermo Fisher Scientific) or 1/5 to 1/2 (v/v) SuperSignal West Femto Maximum Sensitivity Substrate (34095, Thermo Fisher Scientific), chemiluminescent signals were detected using an Amersham™ ImageQuant™ 800 biomolecular imager. Equal loading was validated by staining the PVDF membranes with Ponceau S.

## Acknowledgments

We thank all members of the TSL Support Services, the Center for Plant Molecular Biology (ZMBP) plant cultivation facilities, and Sandra Richter from the ZMBP microscopy platform for their invaluable assistance. We thank Laura Medina-Puch, Delphine Pott, and Shaojun Pan from the ZMBP for generating the mRFP-Lti6b-TurboID construct used in this study.

## Funding

The authors received funding from the Gatsby Charitable Foundation (C.M., A.P., D.L., and S.K.), Biotechnology and Biological Sciences Research Council (BBSRC) BB/P012574 (Plant Health ISP) (S.K.), European Research Council (ERC) 743165 (S.K.), and BASF Plant Science (J.K. and S.K.). D.L. was funded by the DFG Walter Benjamin Programme (project no. 464864389). The funders had no role in the study design, data collection and analysis, decision to publish, or preparation of the manuscript.

## Author contributions

CM: Conceptualization, Formal analysis, Supervision, Validation, Visualisation, Writing – Original draft, Writing – review and editing, Project administration

SK: Conceptualization, Supervision, Project administration, Funding acquisition, Writing – review and editing

HP: Conceptualization, Investigation, Methodology, Validation, Visualisation, Writing – review and editing

AP: Investigation, Writing – review and editing

MPC: Conceptualization, Data Curation, Investigation, Methodology, Validation, Writing – review and editing.

JS: Investigation, Methodology, Writing – review and editing.

DL: Conceptualization, Investigation, Methodology, Writing – review and editing.

JK: Investigation, Resources, Writing – review and editing

## Competing interests

J.K, C.M. and M.P.C. received funding from industry to study NLR biology at the time of the study. S.K. receives funding from industry to study NLR biology and co-founded a start-up company (Resurrect Bio Ltd.). C.M., M.P.C., J.K., and S.K. have filed patents on NLR biology. M.P.C. received fees from Resurrect Bio Ltd.

## Supplementary tables

**Table S1: Constructs generated or used in this study.**

## Supplementary Figures

**Figure S1:**
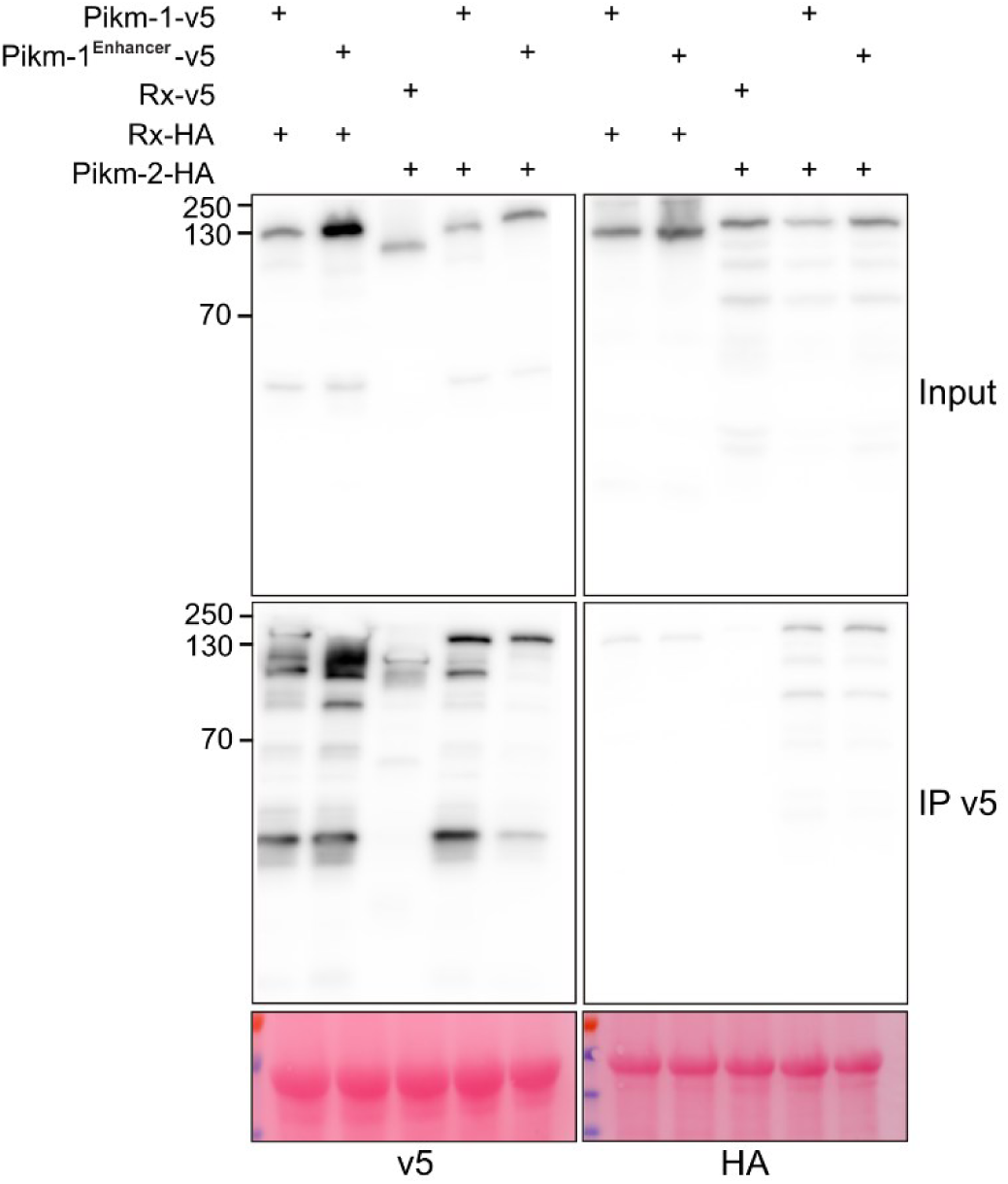
Pikm-1^Enhancer^ interacts with Pikm-2 prior to effector-mediated activation. Replicate of the experiment shown in Figure 1.

**Figure S2.**
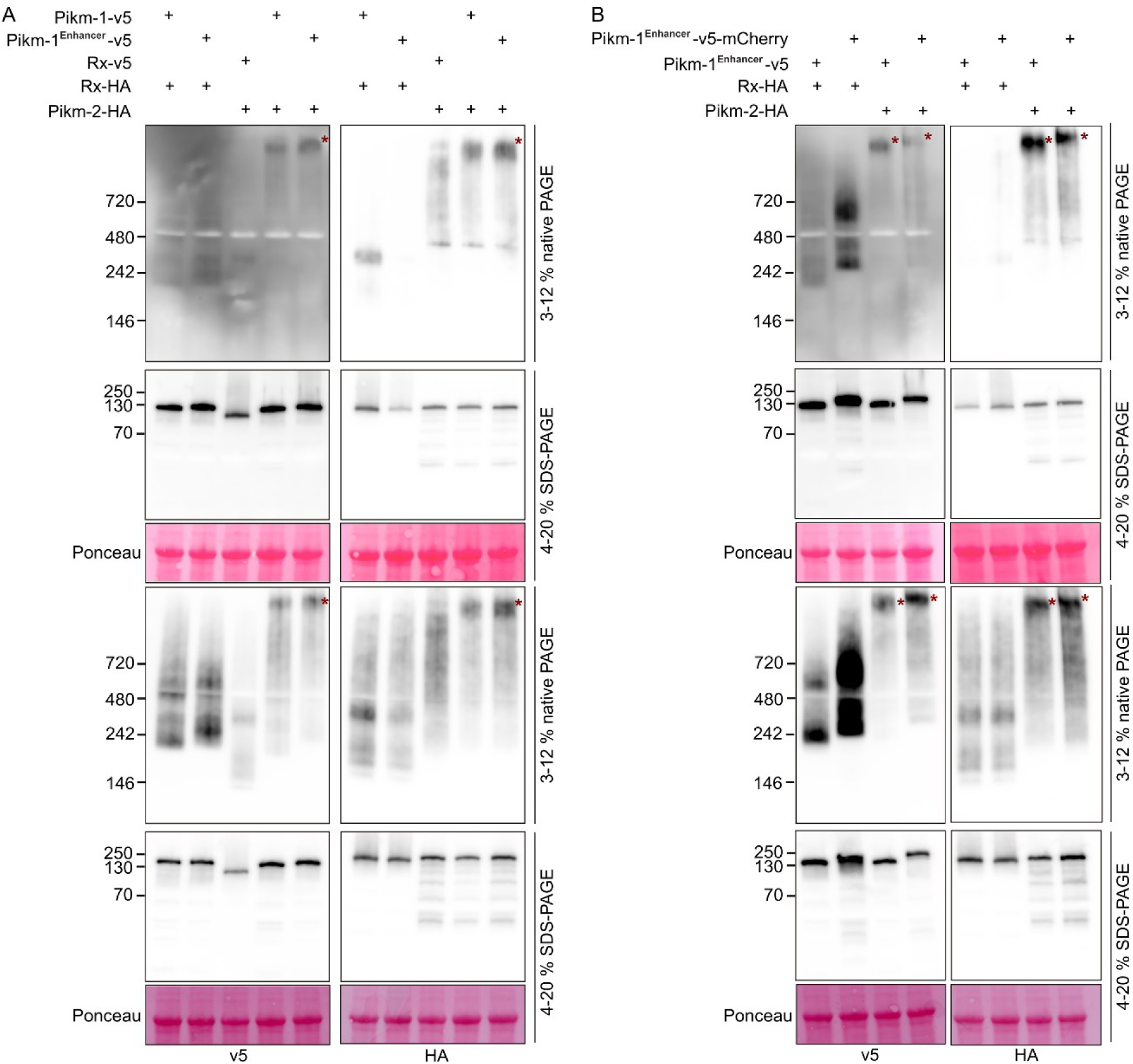
The Pik pair forms a ∼1 MDa hetero-oligomer prior to effector-mediated activation. A-B) Replicates of the experiment shown in Figure 2.

**Figure S3.**
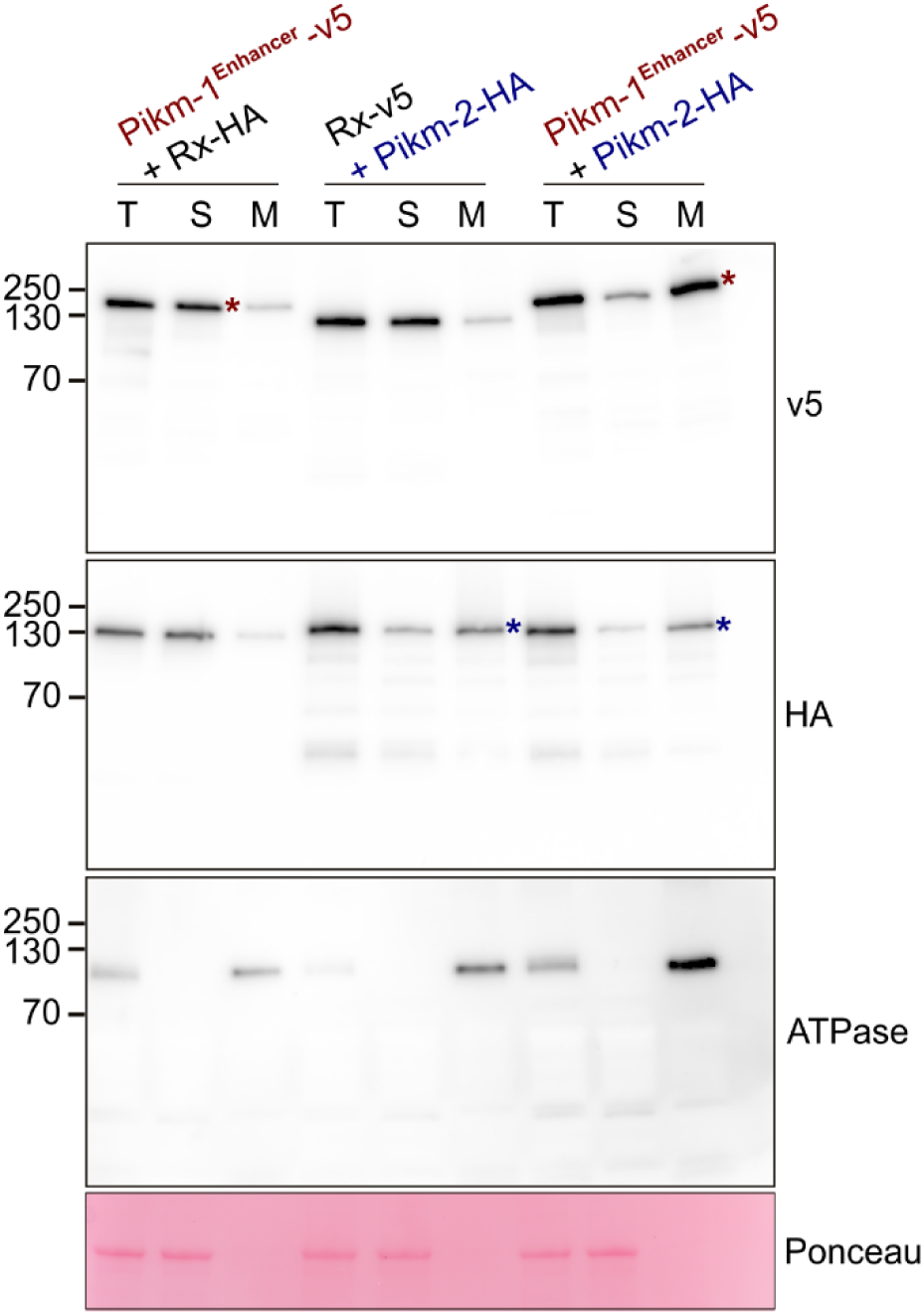
Pikm-1^Enhancer^ and Pikm-2 preferentially accumulate in the membrane fraction when co-expressed. Replicate of the experiment shown in Figure 3.

**Figure S4.**
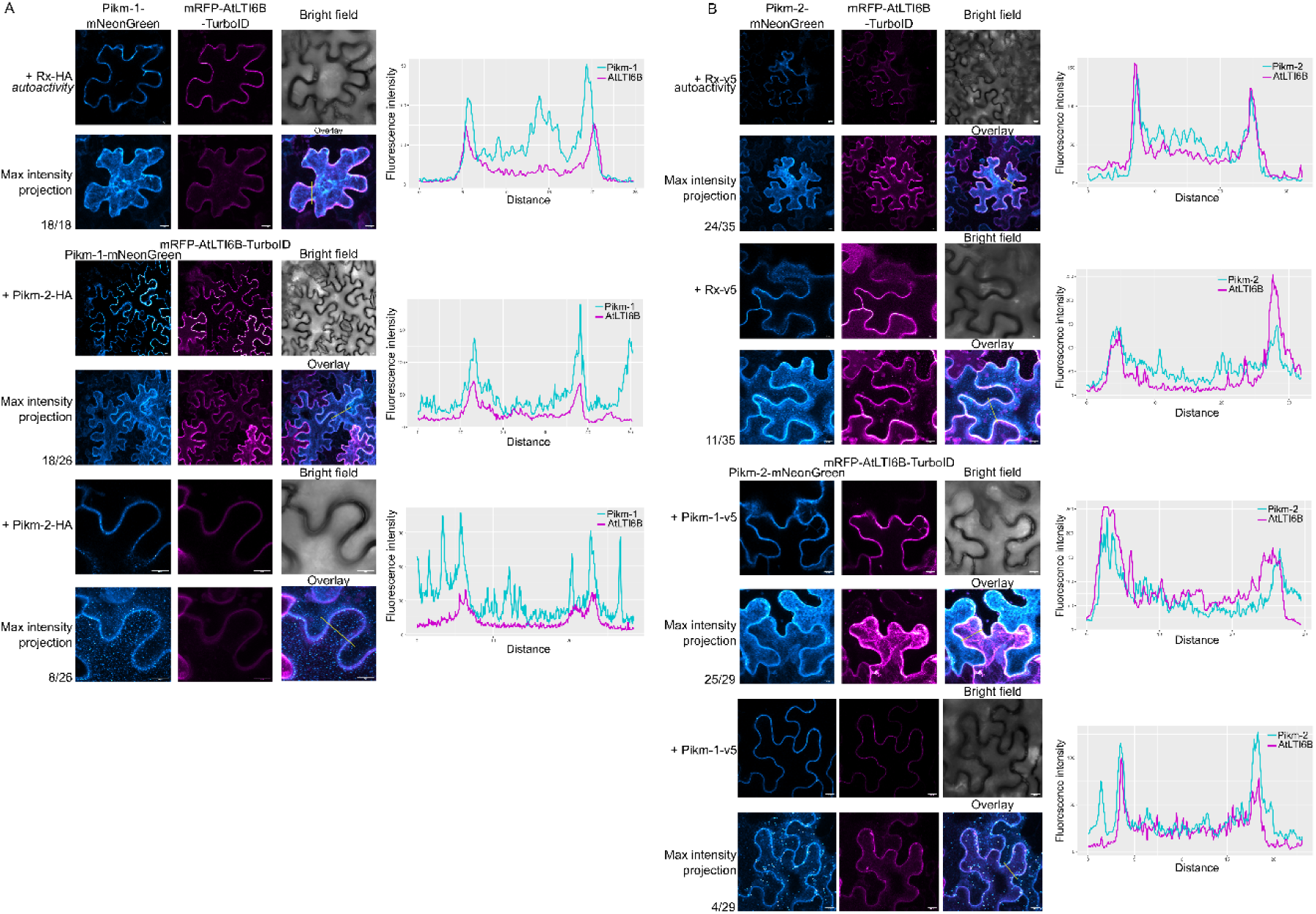
The Pik hetero-complex localises to the plasma membrane. Confocal microscopy images (single plane and maximum projection of *z*-stacks of 10–25 slides) showing the subcellular localisation of A) Pikm-1-v5-mNeonGreen or B) Pikm-2-mNeonGreen co-expressed with Rx or the matching NLR from the Pik pair. LTI6B from Arabidopsis was used as a plasma membrane marker. The graphs at right depict the fluorescence intensity from each channel across the yellow line shown on the overlay image. All components were transiently expressed in *N. benthamiana* via agroinfiltration, and subcellular localisation was observed at 2 dpi. Laser excitation was set to 561 nm (diode) for mRFP and to 488 nm (Argon) for mNeonGreen, and detection wavelengths were set to 590–660 nm and 500–550 nm for mCherry and mNeonGreen, respectively.

**Figure S5.**
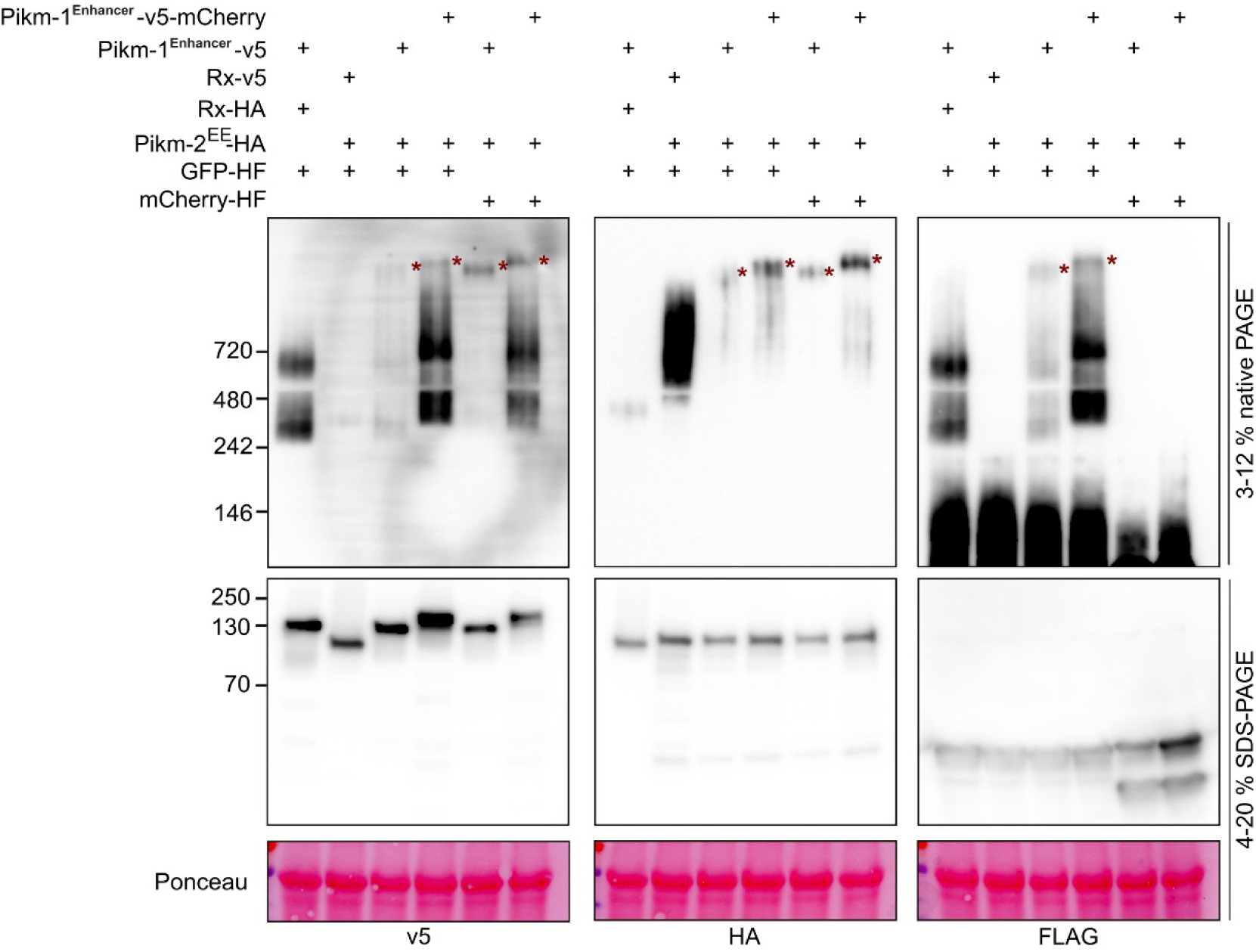
The ligand bound by Pikm-1^Enhancer^ is part of the Pikm-1^Enhancer^/Pikm-2 hetero-oligomer. BN-PAGE (top) and SDS-PAGE (bottom) assays showing that GFP is part of the Pikm-2^EE^ + Pikm-1^Enhancer^ hetero-oligomer and that the corresponding bands shifted when the ‘heavy’ version of Pikm-1^Enhancer^ was used. Co-expression of Rx with Pikm-1^Enhancer^ or Pikm-2^EE^ was used as a negative control for hetero-oligomer formation. mCherry was used as a negative control for the presence of the ligand bound to Pikm-1^Enhancer^. The samples used for BN-PAGE were run in parallel on a denaturing SDS-PAGE gel to ensure all components were expressed. Approximate molecular weights (kDa) of the proteins are shown at left, and red asterisks point to the bands corresponding to the ∼1 MDa Pik oligomer. Ponceau staining showing Rubisco is shown below the immunoblots.

